# Systematic evaluation of 24 extraction and library preparation combinations for metagenomic sequencing of SARS-CoV-2 in saliva

**DOI:** 10.64898/2026.04.16.719115

**Authors:** Kenin Qian, Varada Abhyankar, Dahlia Keo, Payton Zarceno, Traci Toy, Eleazar Eskin, Valerie A. Arboleda

## Abstract

Sequencing the respiratory tract transcriptome has the potential to provide insights into infectious pathogens and the host’s immune response. While DNA-based sequencing is more standard in clinical laboratories due to its stability, RNA assays offer unique advantages. RNA reflects dynamic physiological changes, and for RNA viruses, viral RNA particles directly represent copies of the viral genome, enabling greater diagnostic sensitivity. However, RNA’s susceptibility to degradation remains a significant challenge, particularly in RNase-rich specimens like saliva. To address this, we conducted a systematic, combinatorial evaluation of 24 distinct mNGS workflows, crossing eight nucleic acid extraction methods with three RNA-Seq library preparation protocols. Remnant saliva samples (n = 6) were pooled and spiked with MS2 phage as a control. The SARS-CoV-2 virus was spiked into half of the samples, which were extracted using the eight different extraction methods (n = 3) and compared using RNA Integrity Number equivalent (RINe) scores and RNA concentration. The extracted RNA was then processed across the three library construction methods and subjected to short-read sequencing to assess all 24 combinations head-to-head. We compared methods based on viral read recovery and found that RINe and concentration did not correlate with viral detection. The Zymo Quick-RNA Magbead kit and the Tecan Revelo RNA-Seq High-Sensitivity RNA library kit were the extraction and library-preparation kits that yielded the most SARS-CoV-2 reads, respectively. Importantly, our combinatorial analysis revealed that any small variability attributable to different nucleic acid extraction methods was heavily overshadowed by differences in quality attributable to the RNA-Seq library preparation methods. These findings challenge the reliance on conventional RNA quality metrics for clinical metagenomics and underscore the need to redefine extraction quality standards for mNGS applications.

**IMPORTANCE:** mNGS is a powerful and unbiased approach towards pathogen detection that has mostly been applied to blood and cerebrospinal fluid samples. However mNGS has recently been applied to more areas including the respiratory pathogen detection space, with potential applications in both in-patient diagnostics and public health surveillance. Saliva samples are an ideal sample type for these use cases since they can be collected non-invasively. However, saliva is also a challenging sample type due to its high RNase activity and often yields low-quality nucleic acid. This study explores the feasibility of using saliva specimens in mNGS with contrived SARS-CoV-2 samples to optimize the combination of two factors: nucleic acid extraction and RNA-seq library preparation. Exploration in this area could enhance the sensitivity of saliva-based mNGS assays, with the goal of future expansion of this specimen type in clinical diagnostics and public health surveillance.

**Key Points:**

- The choice of RNA-Seq library preparation kit has a greater impact on pathogen detection than the nucleic acid extraction method.
- The combination of Zymo Quick-RNA Magbead extraction kit and TECAN Revelo RNA-Seq High Sensitivity RNA library kit recovered the highest percentage of total SARS-CoV-2 reads.
- RNA quantity and RINe score do not correlate with viral read capture, indicating a need for an alternative metric to assess RNA quality for downstream mNGS clinical diagnostics.

## INTRODUCTION

The nucleic acids from human tissue interfaces with the environment, capturing valuable information about the human host and colonizing microorganisms. While most microorganisms are commensal, a minority are pathogenic and cause clinical disease. Unbiased sequencing of RNA from clinical specimens offers a powerful approach to 1) identify infectious pathogens and prompt more precision treatments, 2) detect emerging pathogens of pandemic potential, and 3) uncover early outbreaks in healthcare settings. Recent advances, such as the introduction of rapid and cost-effective short-read sequencing-based platforms such as the MiSeq i100, are accelerating the adoption of sequencing as a frontline diagnostic tool for infectious disease, mirroring its widespread use in cancer^1^, newborn screening^2^, and pediatric germline testing^3,4^.

DNA is routinely sequenced in inherited and somatic tumor mutation testing, as DNA encodes all possible RNA transcripts and proteins. However, despite being less stable than DNA^5^, RNA has several advantages over DNA. First, mRNA represents a functional output of the genome after splicing, providing direct evidence of translational consequences^6^. Second, RNA molecules exist in higher copy numbers compared to corresponding DNA molecules, improving the assay’s sensitivity and highlighting actively expressed transcripts in a given tissue^7,8^. Third, RNA sequencing enables the detection of a broad range of pathogens, including RNA viruses, DNA viruses, fungi, and bacterial pathogens, all of which utilize RNA to direct protein translation. While RNA is already well-established as a target in PCR-based viral diagnostics, non-targeted approaches such as metatranscriptomic sequencing are believed to require high-quality RNA for efficient cDNA synthesis and sufficient pathogen read abundance for accurate detection. Challenges to RNA-based diagnostics remain, particularly due to RNA’s susceptibility to degradation and its tissue-, cell-, and allele-specific expression patterns, all of which contribute to variability in sequencing results. Nonetheless, overcoming these obstacles could unlock substantial improvements in the sensitivity and specificity for detection of a wide range of viral pathogens.

The scale of diagnostic testing during the COVID-19 pandemic significantly increased public and clinical familiarity with self-collection of upper respiratory and oral specimens. It accelerated the development of methods to extract high-quality nucleic acids suitable for sequencing-based readouts. While these early approaches relied on targeted PCR for a limited number of pathogens^9,10^, extending to an unbiased metagenomic sequencing requires a reassessment of the minimal RNA quality and quantity thresholds necessary for accurate RNA-Seq-based diagnostics.

In this study, we evaluated a combination of eight nucleic acid extraction kits, either DNA/RNA or RNA only, and three RNA-Seq library preparation kits. We assessed the quality of the extracted nucleic acid samples, the suitability of library construction, and the diagnostic sensitivity of each extraction–library prep combination in an mNGS workflow. We tested kits that fulfilled several key criteria for scalability (throughput), reliability, and speed of extraction for saliva samples. Extraction performance was assessed using standard measures such as RNA Integrity Number equivalent (RINe) score and total RNA yield. Additionally, we created RNA-Seq libraries from extracted nucleic acid to compare the extraction methods based on RNA counts from downstream RNA library sequencing results and to compare the different RNA-Seq library preparation methods in their application to mNGS. We found that the MagMAX mirVana Total RNA and the Zymo Quick-RNA Magbead RNA extraction kits preserve the most spiked-in viral reads in our contrived saliva samples. However, overall, the choice of RNA-Seq library preparation kit had a larger impact on viral read preservation, coverage and distribution of reads across the SARS-CoV-2 genome. We found that the Revelo RNA-Seq High Sensitivity library preparation kit, in combination with the Zymo Quick-RNA Magbead RNA extraction kit, yielded the highest percentage of spiked-in SARS-CoV-2 reads. Our study showed no correlation between nucleic acid extraction quantity or RINe score and preservation of viral reads, suggesting the need for alternative quality metrics when assessing RNA samples in an mNGS context.

## METHODS

### Creating Contrived Saliva Samples

We generated contrived negative saliva samples by pooling 96 SARS-CoV-2 PCR negative saliva samples from healthy individuals and adding MS2 phage (Varizymes 6000L) with a final concentration of 12,500 copies/μL. Negative sample matrix were de-identified remnant samples from the UCLA SwabSeq laboratory (#IRB-24-0270) that tested negative for SARS-CoV-2 accessed October 2024.

We generated contrived positive saliva samples by pooling 96 SARS-CoV-2 PCR-negative saliva samples from healthy individuals, adding MS2 phage (Varizymes, 6000L) with a final concentration of 12,500 copies/μL, and adding Heat-inactivated SARS-CoV-2 (ATCC, VR-1986HK) with a final concentration of 5,000 copies/μL. Three positive and three negative samples were made, aliquoted for the 8 different nucleic acid extraction methods, and frozen at -80°C until use. Each aliquot underwent a single freeze-thaw cycle before extraction.

### Total Nucleic Acid Extraction from Saliva Samples

We used the following kits all following manufacturer protocol: Zymo Quick-RNA MagBead (Zymo Research, R2132), Zymo Quick-DNA/RNA Viral MagBead (Zymo Research, R2141), MagMAX mirVana Total RNA Isolation Kit (ThermoFisher, A27828), MagMAX Viral/Pathogen Nucleic Acid Isolation Kit (ThermoFisher, A42352), MagMAX Microbiome Ultra Nucleic Acid Isolation Kit (ThermoFisher, A42357), MagMAX-96 Viral RNA Isolation Kit, (ThermoFisher, AM1836), QIAamp Viral RNA Mini Kit (Qiagen, 52904), and PureLink Viral RNA/DNA Mini Kit (Fisher Scientific, 12280050). Each extraction was done in triplicate for the positive and negative samples. The input volume of unextracted saliva was 200 μL, and the purified nucleic acid was eluted in 50 μL of elution buffer. Extractions were performed manually by a single operator using identical pipettes and equipment.

### TapeStation RNA Quality Control

Purified RNA samples were analyzed on the 4150 TapeStation System (Agilent, G2992AA) using the Agilent TapeStation Analysis Software version 4.1.1. All RNA was done on High Sensitivity RNA gels and buffers. 1 μL of sample RNA was added to 2 μL of High Sensitivity RNA Sample Buffer, vortexed for 1 minute, and then incubated for 3 minutes at 72°C. The sample was placed on ice for 2 minutes, quickly spun down, and subsequently analyzed.

### Qubit Nucleic Acid Quantification

Purified RNA samples were analyzed on the Qubit 4 Fluorometer (ThermoFisher, Q33238) using the Qubit RNA High Sensitivity Assay Kit (ThermoFisher, Q32852).

### RNA Library Generation and Sequencing

We used the High Sensitivity BRB-Seq kit (Alithea Genomics, #11881) following the manufacturer’s protocol to generate the BRB-Seq libraries. We used the NEBNext Single Cell/Low Input RNA Library Prep Kit for Illumina (New England Biolabs, E6420L) along with the NEBNext Multiplex Oligos for Illumina (New England Biolabs, E6440L) according to the manufacturer’s protocol to generate NEB libraries. We used the Revelo RNA-Seq High Sensitivity library preparation kit (TECAN, 30201359) with SPIABoost Human RMG (TECAN, 30201372), and the TECAN Adaptor Index Plate (TECAN, 30219997) following manufacturer protocol to generate TECAN libraries. All libraries were generated from 10 ng of RNA when available. When RNA was unquantifiable, the maximum volume of input RNA was added. The library was run on TapeStation to ensure quality and check for excess primer dimer and adaptor dimer formation. We used the NextSeq 2000 (Illumina, 20038897) and P2 100-cycle flow cells (Illumina, 20044468) for sequencing RNA libraries. The cycles for the BRB-Seq libraries were 28 for read 1, 74 for read 2, 8 for index 1 and 2, 51 for read 1 and 2, and 8 for index 1 and 2 for the NEB libraries, and 51 for read 1 and 2, and 10 for index 1 and 2 for the TECAN libraries. The reads were demultiplexed using the Illumina bcl2fastq software for the generation of FASTQ files.

### Data Analysis

The FASTQs’ quality was assessed using FastQC^11^, and the reads were classified using the Kraken 2 classifier^12^. FASTQ files were aligned to the SARS-CoV-2 genome using Refseq^13^ GCF_009858895.2_ASM985889v3 with STAR aligner^14^ using local alignment, restricting mismatches per read to 2, requiring a minimum alignment score of 90, and the removal of multimapped reads. FASTQ files can be accessed from SRA with accession number PRJNA1401922.

## RESULTS

### Generation of study specimens for nucleic acid extraction testing

We generated contrived specimens to directly compare the eight nucleic acid extraction methods across quality and quantity metrics. To prevent individual specimen differences from biasing our study of extraction quality, we created a pooled source of remnant saliva specimens from 96 individuals who had tested negative on a SARS-CoV-2 Swabseq test^9^ to create a uniform background in which we could add specified quantities of SARS-CoV-2. As an internal control for nucleic acid extraction, we added bacteriophage MS2, which is commonly used to control the quality of nucleic acid extraction in clinical RT-PCR-based assays^15^ to a final concentration of 12,500 copies/μL. In our sequencing readout, quantification of MS2 phage reads serves as a comparative measure of overall extraction efficacy across the different extraction methods. Samples were then split into a positive and negative pool, and the SARS-CoV-2 heat-inactivated virus was added to the positive pool at a concentration of 5,000 copies/μL (**Figure 1A**). This moderate-to-high concentration was selected to ensure that copies of SARS-CoV-2 would be detectable, based on our previous limit-of-detection studies^9^ for the SwabSeq assay and to allow us to compare the ability of each extraction method to retain viral RNAs. In total, we created eight identical sets of contrived positive (n=3) and negative (n=3) saliva samples, one set for each tested nucleic acid extraction kit (**Table 1**).

**Figure 1.**
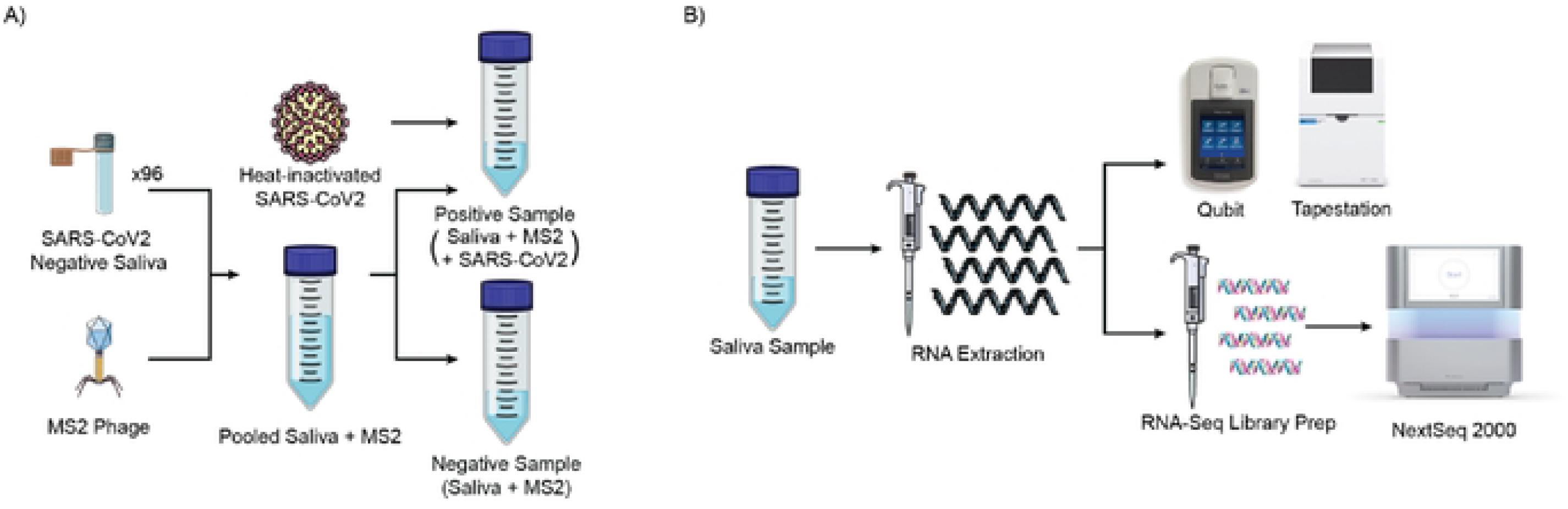
Study Design. Positive and negative contrived saliva samples were created to test eight nucleic acid extraction methods for mNGS RNA testing. A) SARS-CoV-2 negative saliva samples were collected and pooled from 96 individuals. MS2 phage was added to the pool at a final concentration of 12,500 copies/μL. The pooled saliva and MS2 phage samples were split into positive and negative samples. Heat-inactivated SARS-CoV-2 was added to the positive sample with a final concentration of 5,000 copies/μL. B) Saliva samples were then extracted using one of eight different extraction kits, and RNA-Seq libraries were generated with one of three library preparation kits. The extracted nucleic acid was assessed for quality and quantity using TapeStation and Qubit. RNA-Seq libraries were then sequenced, and the SARS-CoV-2 and MS2 reads were quantified.

**Table 1.** Comparison of nucleic acid extraction kits by method and reagents. 2-mercaptoethanol (2-BME), Isopropyl Alcohol (IPA), Proteinase K (PK).

| Extraction Kit Comparison |  |  |  |  |  |  |  |  |  |
| --- | --- | --- | --- | --- | --- | --- | --- | --- | --- |
| Kit | Abbr. | Nucleic Acid Type | Extraction Method | 2-BME | IPA | Carrier RNA | PK | DNase | Automation Compatibility |
| Zymo Quick-RNA Magbead | Z1 | DNA/RNA | Magnetic Beads | No | Yes | No | Yes | No | KingFisher |
| Zymo Quick-DNA/RNA Viral Magbead | Z2 | DNA/RNA | Magnetic Beads | Yes | Yes | No | Yes | No | KingFisher |
| MagMAX mirVana Total RNA | M1 | RNA | Magnetic Beads | Yes | Yes | No | Yes | Yes | KingFisher |
| MagMAX Viral/Pathogen | M2 | DNA/RNA | Magnetic Beads | No | No | No | Yes | No | KingFisher |
| MagMAX Microbiome Ultra | M3 | DNA/RNA | Magnetic Beads | No | No | No | Yes | No | KingFisher |
| MagMAX Viral RNA Isolation | M4 | DNA/RNA | Magnetic Beads | No | Yes | Yes | No | No | KingFisher |
| QIAamp Viral RNA | Q1 | DNA/RNA | Column Filter | No | No | Yes | No | No | QiaCube |
| PureLink Viral RNA/DNA | P1 | DNA/RNA | Column Filter | No | No | Yes | Yes | No | N/A |

### Comparing nucleic acid extraction quality from eight nucleic acid extraction kits

In the clinical setting, oropharyngeal and respiratory specimens are collected to identify the causal pathogen in respiratory illnesses. However, these samples are typically self-collected in a manner that may not have preservatives to prevent RNA degradation that naturally occurs due to heat and endogenous RNases. We tested eight nucleic acid extraction methods in saliva samples, each with slight variations in techniques and reagents, to directly compare and maximize the quality of the extracted nucleic acid (**Table 1**). These extraction kits are sourced from three independent suppliers and can be semi-automated on the Kingfisher Apex (Thermofisher) or QIAcube Connect (Qiagen) platforms. They are designed to extract total nucleic acids (DNA and RNA) or only RNA.

We opted to test both total nucleic acids and RNA-only based extraction methods as we felt that the presence of DNA could potentially improve enrichment of DNA-based viruses such as adenovirus^16,17^. Furthermore, the first step in the RNA-Seq library preparation is the generation of cDNA from the single-stranded RNA fragments. The presence of genomic DNA (from both human and microbiology organisms) could potentially allow for extension of this test into non-RNA respiratory viruses, although this is not explicitly tested here. We compared each nucleic acid extraction kit by concentration of extracted nucleic acid, RNA Integrity Score (RINe) score, SARS-CoV-2 RNA counts, and MS2 phage RNA counts (**Figure 1B**). Then, for each kit we used three different RNA-Seq library preparation kits (**Table 2**), allowing for comparison of the various RNA-Seq library preparation methods in combination with the nucleic acid extraction kits. Our data provide insight into optimizing both specimen extraction and RNA-Seq library generation for unbiased metagenomic virus detection.

**Table 2.** Comparison of RNA-Seq preparation kits by coverage, priming method, amplification method, and approximate turnaround time (Approx. TAT). Single Primer Isothermal Amplification (SPIA), Polymerase Chain Reaction (PCR).

| RNA-seq Library Preparation Kit | Coverage | Priming Method | Amplification Method | Approx. TAT |
| --- | --- | --- | --- | --- |
| NEBNext Single Cell/Low Input RNA Library Preparation Kit | Full-length | Poly-A and Template Switching | PCR | 8 hours |
| Revelo RNA-Seq High Sensitivity Library Preparation Kit | Full-length | Poly-A and Random | SPIA | 6 hours |
| High Sensitivity BRB-Seq RNA Library Preparation Kit | 3' End | Poly-A | PCR | 4 hours |

**Table 2.** Comparison of RNA-Seq preparation kits by coverage, priming method, amplification method, and approximate turnaround time (Approx. TAT).
| RNA-Seq Library Preparation Kit Comparison |  |  |  |  |
| --- | --- | --- | --- | --- |
| RNA-seq Library Preparation Kit | Coverage | Priming Method | Amplification Method | Approx. TAT |
| NEBNext Single Cell/Low Input RNA Library Preparation Kit | Full-length | Poly-A and Template Switching | PCR | 8 hours |
| Revelo RNA-Seq High Sensitivity Library Preparation Kit | Full-length | Poly-A and Random | SPIA | 6 hours |
| High Sensitivity BRB-Seq RNA Library Preparation Kit | 3' End | Poly-A | PCR | 4 hours |

### The concentration and RINe score of extracted nucleic acid are not correlated with the detection of SARS-CoV-2 reads by sequencing

Nucleic acid concentration is a fundamental quality-control metric for assessing nucleic acid extraction performance. If a method can extract more RNA from the same amount of input, it should better reflect the RNA distribution of the input. Since every extraction kit used identical input specimen volumes, and we adjusted the methods for identical final elution volumes, we could make direct comparisons by the eluted RNA’s concentration. The DNA/RNA and RNA-only extractions were quantified using the Qubit RNA High Sensitivity Assay, which selectively quantifies RNA and is not affected by the presence of DNA. We found that the MagMAX Viral/Pathogen kit and the PureLink Viral RNA/DNA kit had the highest concentration of extracted nucleic acid in both SARS-CoV-2-positive and negative samples. The Zymo Quick-DNA/RNA Viral Magbead kit was the only extraction kit that failed to produce detectable RNA via Qubit RNA High Sensitivity Assay Kit in all three positive and negative extracted nucleic acid replicates. The Zymo Quick-RNA Magbead kit yielded the lowest detectable concentration, and the remaining kits performed comparably **(Table S1)**, all with concentrations around 10 ng/μL, as determined by RNA concentration (**Figure 2A**).

**Figure 2.**
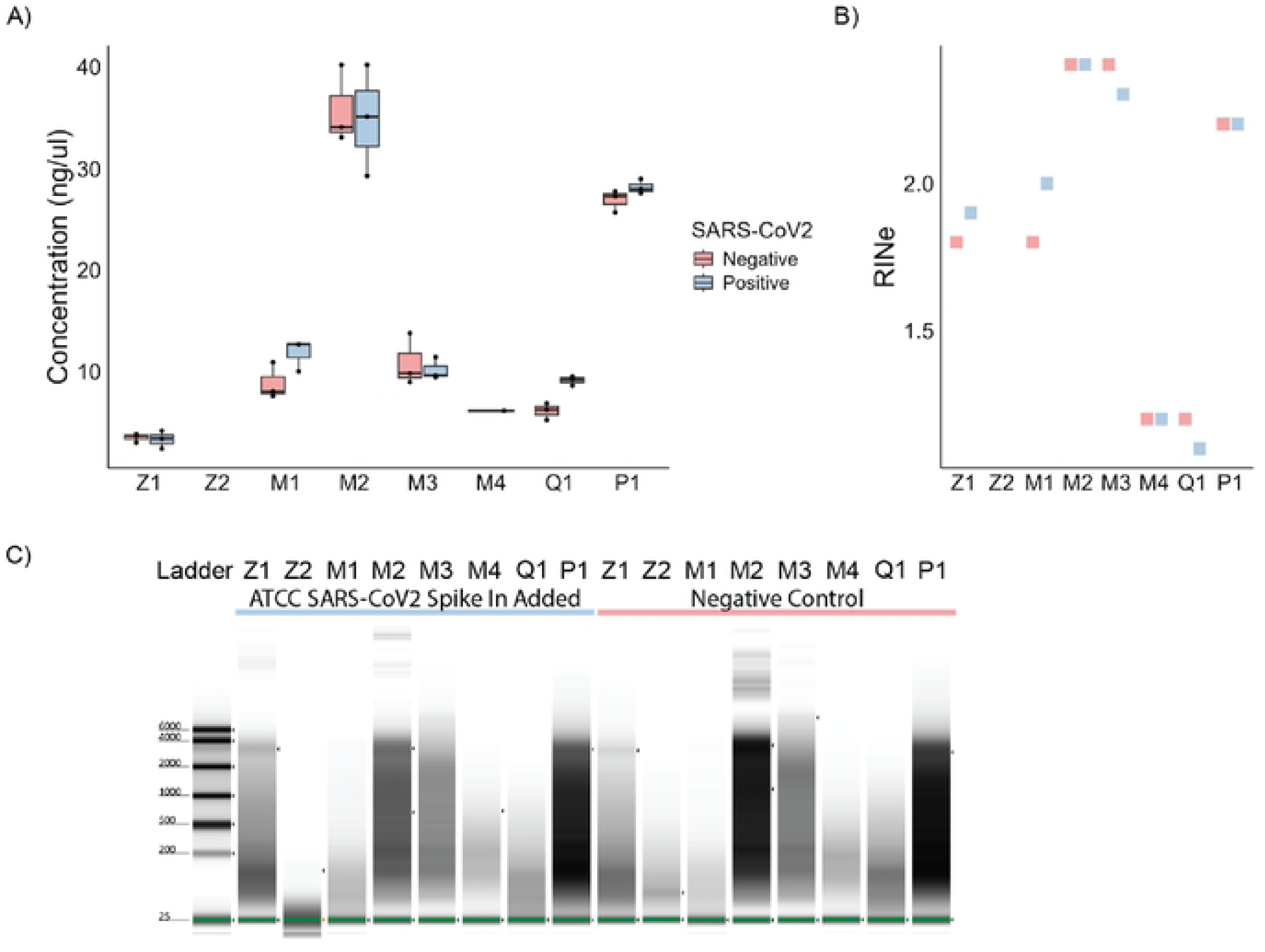
Standard nucleic acid extraction quality metrics show a range of RNA concentrations and universally low RINe scores across all extraction methods. RNA from eight nucleic acid extraction kits were assessed based on A) concentration (ng/μL), B) RINe score, and C) representative tapestation RNA gel electrophoresis. (Z1 - Zymo Quick-RNA Magbead, Z2 - Zymo Quick-DNA/RNA Viral Magbead, M1 - MagMAX mirVana Total RNA, M2 - MagMAX Viral/Pathogen, M3 - MagMAX Microbiome Ultra, M4 - MagMAX Viral RNA Isolation, Q1 - QIAamp Viral RNA, P1 - PureLink Viral RNA/DNA)

The second standard metric used to assess RNA quality for sequencing assays is the RINe score from the Agilent TapeStation, which utilizes microfluidics-based technology to analyze DNA or RNA^18^. RINe is scored on a range from 0-10, where 10 is high-quality RNA showing two ribosomal RNA (rRNA) bands at 1800 and 4000 nucleotides, while a score less than 3 represents RNA without two rRNA bands, indicating degraded RNA. Compared to previous methods, the RINe score assesses the entire electrophoresis trace, rather than just the ratio of the two rRNA bands. Across all our extraction protocols, none of the RINe scores exceeded 3; this is far below the recommended score of 7 for RNA-Seq library protocols. Once again, the MagMAX Viral/Pathogen and PureLink Viral RNA/DNA kits were among the highest scorers, along with the MagMAX Microbiome Ultra kit. The difference between the higher RINe scores and the lower RINe scores was much smaller than with the concentrations. Again, the Zymo Quick-DNA/RNA Viral Magbead was the only kit that failed to provide a RINe score **(Figure 2B)**. The RNA gel shows differences in RNA concentration and RINe score across the extraction methods **(Figure 2C)**.

### RNA-Seq library preparation kits can make up for poor RNA quality

Despite the low RINe scores, we found that nearly all the extraction kits could create RNA-Seq libraries, using three different library preparation methods, with measurable levels of the spiked-in SARS-CoV-2 and MS2. The RNA-Seq library preparation methods were selected for their high sensitivity to detect low-abundance of RNA. The three kits represent a range of turnaround times for library creation **(Table 2)**. At the start of RNA-Seq library preparation, we used equal starting concentrations of extracted nucleic acid, except for the Zymo Quick-DNA/RNA Viral Magbead kit where the maximum volume suggested per kit was added due to the undetectable RNA concentration. Each library had no significant difference in total number of classified reads (**Figure S1**). Given the equal starting conditions for the RNA-Seq libraries, we used the percentage of reads mapping to either SARS-CoV-2 or to MS2 to compare both the different extraction methods for their ability to preserve non-human reads through the extraction process and the different RNA-Seq library preparation methods for their sensitivity to metagenomic sequences.

We first examined the percentage of reads mapping to SARS-CoV-2, grouped by nucleic acid extraction method. The two extraction kits that preserved the highest number of SARS-CoV-2 reads **(Table S3)** were the Zymo Quick-RNA Magbead kit and the MagMAX mirVana Total RNA kit **(Figure 3A, Table S2)**. When grouping the SARS-CoV-2 read percentage by RNA-Seq library preparation kit we found that the Revelo kit returned the highest number of SARS-CoV-2 reads and the BRB-Seq kit returned the lowest recovered SARS-CoV-2 reads **(Figure 3B, Table S3)**. The spiked-in MS2 also showed the MagMAX mirVana Total RNA kit has the highest average MS2 read percentage. Interestingly, the Zymo Quick-RNA Magbead kit’s MS2 read percentage appeared to be dependent on the RNA-Seq library preparation kit used rather than extraction method **(Figure 3C).** When comparing RNA-Seq library preparation kits, the spiked-in MS2 was consistently detected with the Revelo and NEB kits. The BRB-Seq kit was unable to detect MS2 in the majority of the extracted nucleic acid kits, likely due to the kits’ use of oligo-dT primers which are incompatible with MS2 RNA priming, as a bacteriophage MS2 does not contain a poly-A tail and any capture is likely due to an adenine-rich region of the MS2 genome. The Revelo library preparation kit yielded the highest number of spiked-in reads (MS2 and SARS-CoV-2) across all kits (**Figure 3D**) and in combination with Zymo Quick-RNA Magbead RNA **(Figure 3E, 3F)**.

**Figure 3.**
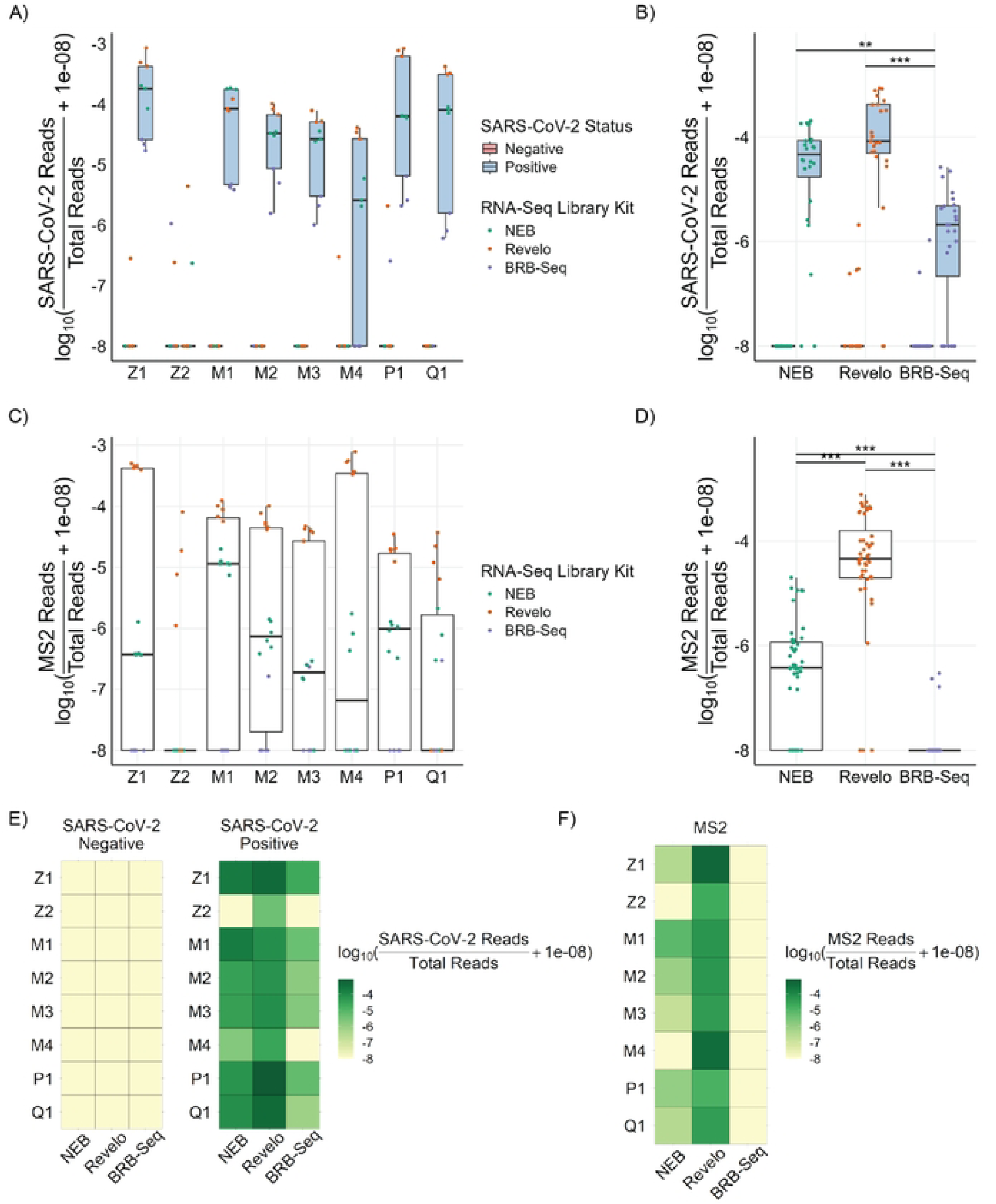
RNA-Seq library read quantification detects MS2 and SARS-CoV-2 across nearly all extraction methods. RNA-Seq libraries were created from the eight different nucleic acid extraction methods using three different RNA-Seq library preparation methods: NEBNext Single Cell/Low Input RNA library preparation kit (NEB), Revelo RNA-Seq High Sensitivity library preparation kit (Revelo), and High Sensitivity BRB-Seq RNA library preparation kit (BRB-Seq). Reads were mapped to viral, bacterial, and eukaryotic genomes present in the RefSeq database. The proportion of total reads mapping to SARS-CoV-2 is shown either grouped by A) nucleic extraction method or B) RNA-Seq library preparation method. As a measure of extraction quality, the proportion of total reads mapping to MS2 is shown either grouped by C) nucleic extraction method or D) by RNA-Seq library preparation method. The proportion of E) SARS-CoV-2 reads or F) MS2 reads to total reads is shown with every combination of nucleic extraction methods and RNA-Seq library preparation kits with a darker shade indicating a higher proportion of SARS-CoV-2 or MS2 reads. (Z1 - Zymo Quick-RNA Magbead, Z2 - Zymo Quick-DNA/RNA Viral Magbead, M1 - MagMAX mirVana Total RNA, M2 - MagMAX Viral/Pathogen, M3 - MagMAX Microbiome Ultra, M4 - MagMAX Viral RNA Isolation, Q1 - QIAamp Viral RNA, P1 - PureLink Viral RNA/DNA). Statistics were calculated by a One-Way ANOVA, followed by Tukey’s HSD post-hoc test. Asterisks above connecting brackets indicate significant differences between those two specific groups. * denotes p<0.05, ** denotes p<0.01, and *** denotes p<0.001.

**Figure 4.**
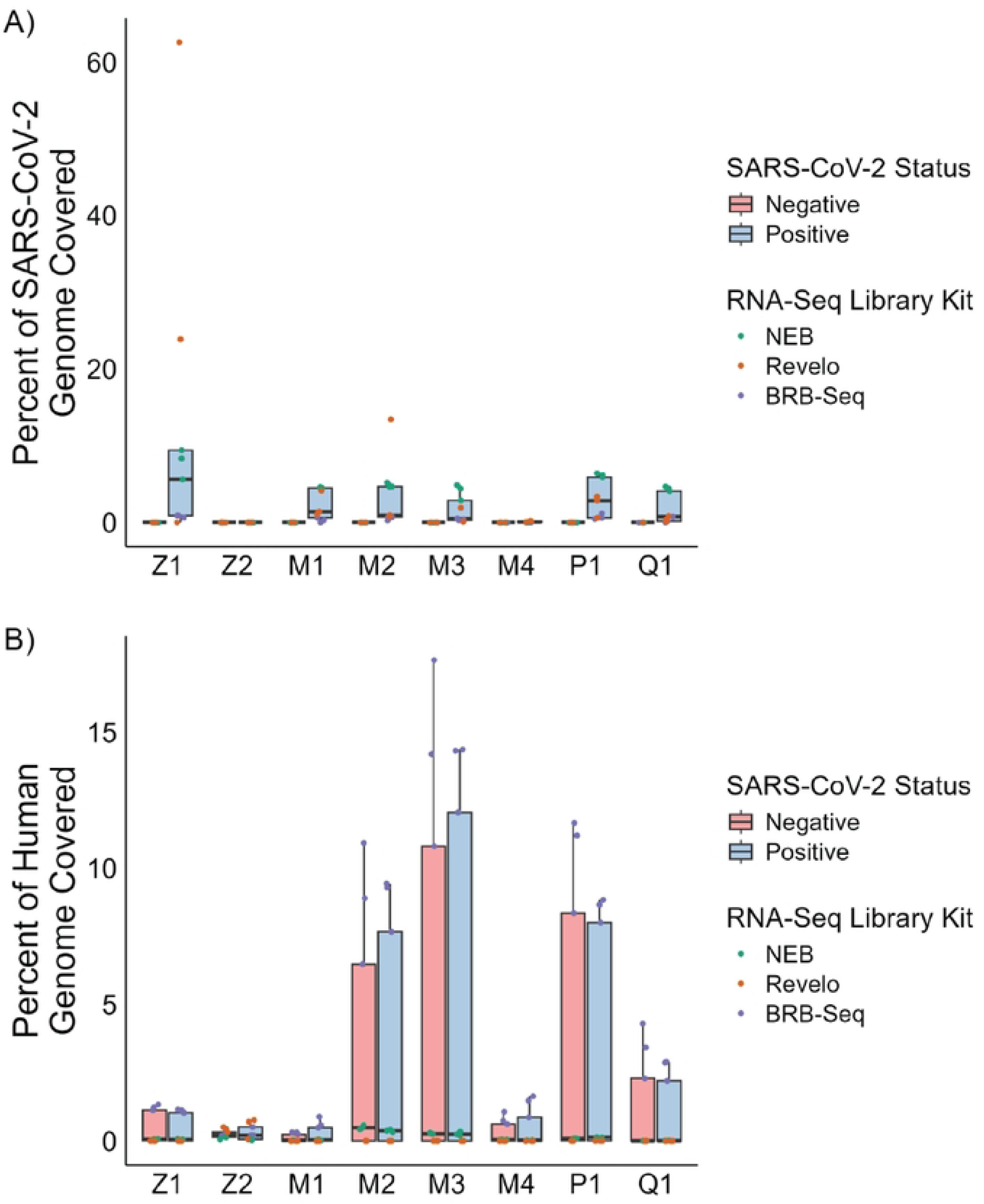
Genome coverage is low for both SARS-CoV-2 and Human genomes across all extraction and RNA-seq library preparation methods. Reads from RNA-seq libraries made from RNA extracted from saliva samples mapped against A) SARS-CoV-2 genome or B) Human genome to show genome coverage with a minimum depth of five reads. (21 - Zymo Quick-RNA Magbead, Z2 - Zymo Quick-DNNRNA Viral Magbead, M1 - MagMAX mirVana Total RNA. M2 - MagMAX Viral/Pathogen, M3 - MagMAX Microbiome Ultra, M4 - MagMAX Viral RNA Isolation, Q1 - QIAamp Viral RNA, P1 - Purelink Viral RNA/DNA)

A unique aspect of RNA-seq library preparation methods is that in the sequencing-based approach, the overwhelming majority of the classified sequencing reads do not map to the pathogen of interest and instead map to the host genome and transcriptome and the microbiome. This is particularly prominent in non-sterile specimens, such as saliva. In clinical diagnostic mNGS assays, the percent of reads mapping to the pathogen genome detected is typically less than 1% of total classified reads and the number of total classified reads can vary widely^19^. In our study, we find that the proportion of reads uniquely mapping to SARS-CoV-2 was low (Median: 2.89x10^-5^ IQR: 2.09x10^-6^- 8.47x10^-5^), but significantly higher than negative samples in which 66/72 samples had 0 classified reads mapping to SARS Co-V2 and the other 6 samples had between 1-6 reads (Median: 0 IQR: 0 - 0).

We next asked whether there was a relationship between extraction measures, such as concentration and RINe score, and sequencing-based measures of library quality. There was no appreciable relationship between RNA concentration or RINe score with either SARS-CoV-2 reads or MS2 reads (**Figure S2)**. Moreover, we observed that the percentage of duplicate reads in the RNA-Seq libraries remained relatively consistent across the RNA extraction methods, but was more affected by the different library preparation kits (**Figure S3, Table S4**).

Finally we assessed the genomic distribution of reads in our sequenced libraries. Classification of reads showed a majority of reads mapped to non-viral sources. Eukaryotic reads dominated in RNA-seq library preparation methods without rRNA depletion: BRB-Seq and NEB. The Revelo kit includes an intrinsic rRNA depletion step that increases the bacterial read percentage (**Figure S4**). Viral genome mapping reveals that SARS-CoV-2 reads, although sparse, are distributed across the SARS-CoV-2 genome rather than originating from only one or two locations, with the breadth of this distribution depending on the RNA-seq library preparation method. The two oligo-dT based RNA-seq library preparation methods, NEB and BRB-Seq, showed increased SARS-CoV-2 coverage at the 3’ end, consistent with highly fragmented RNA. In contrast, the Revelo libraries showed relatively stable coverage of the SARS-CoV-2 genome due to the use of both oligo-dT and random primers in the library preparation (**Figure S5**). Quantification of the coverage from our libraries to the SARS-CoV-2 genomes had an IQR from 0.7% - 4.3% in the SARS-CoV-2 positive samples and an IQR of 0% in the SARS-CoV-2 negative samples (**Figure S6**).

## DISCUSSION

We evaluated eight nucleic acid extraction kits and three RNA-Seq library preparation methods, comparing them based on their ability to detect SARS-CoV-2 in an unbiased metagenomic sequencing assay. This project serves as a proof-of-principle, that even from degraded, respiratory specimens, we can reliably detect spiked-in SARS-CoV-2. This technology can be extended to other respiratory pathogens, however, to what extent this would be feasible and scalable has not been published. To compare each extraction method, both separately and in combination with three different library preparation kits, we generated contrived samples that were either negative or positive for SARS-CoV-2. These enabled us to directly compare the efficacy of extraction and library preparation across the multiple replicates. In total, 24 combinations were tested, with 3 SARS-CoV-2 and 3 negative samples for each combination.

This is the first time an assessment of multiple combinations of RNA extraction kits and library preparation methods has been performed to evaluate recovery of viral pathogens, essentially mimicking samples with a single pathogen infection. Our work demonstrated some variability between nucleic acid extraction kits, as assessed by RINe score and RNA concentration. However, all nucleic acid extractions yielded degraded and low levels of nucleic acid. Despite this variability, RNA-Seq libraries generated from the extracted nucleic acid successfully recovered SARS-CoV-2 and MS2 from all but one nucleic acid extraction method.

When comparing RNA-Seq library preparation methods, we found that BRB-Seq was the fastest and, at this time, the least costly, and could still detect SARS-CoV-2; however, it had the lowest recovered spiked-in read percentage, missing MS2 in all extracted nucleic acid methods, highlighting the limitations of 3’ only priming as the MS2 bacteriophage lacks poly-A tails. Revelo yielded the highest recovered spiked-in read percentage but also produced libraries with the highest duplicated-read rate. In contrast, NEB could recover SARS-CoV-2 and MS2 while producing libraries with the lower percentage of duplicated read rate. We observed that both oligo-dT priming methods were heavily biased towards the 3’ end of the SARS-CoV-2 genome. While this was expected, given that BRB-Seq is designed to sequence the 3’ end and the highly degraded sample quality prevented NEB from capturing many fully intact SARS-CoV-2 RNA molecules, it does highlight a limitation of isolated oligo-dT based approaches, especially when working with low-quality RNA. The Revelo kit contained oligo-dT primers but was also supplemented with random hexamer priming that allowed for greater coverage of the SARS-CoV-2 genome. Additionally, the Revelo uses enzymatic shearing followed by end-repair and A-tailing which also improves oligo-dT based coverage. Overall, all three library preparation methods provided enough genome coverage to confidently detect SARS-CoV-2, but differences in the molecular technology changed the number of classified reads across the pathogen genome coverage and likely choice of kit will affect the sensitivity towards pathogen detection.

### Scalability versus Accuracy

One of the benefits of sequencing-based assays is the potential for increasing scale without requiring additional machines. While RT-PCR-based diagnostic assays require more machines to double or triple throughput, our team has leveraged automated workflows, coupled with sequencing, to rapidly scale from hundreds to thousands of samples per day without requiring significant changes to capacity. All of the extraction kits tested, except for the PureLink Viral RNA/DNA kit, were compatible with automated platforms such as the KingFisher series of machines commonly used in clinical laboratories. RNA-Seq-based library prep platforms are significantly more complicated relative to our targeted PCR platforms^9,20^. While full-transcriptome library preparation kits have increased the number of reads for SARS-CoV-2, they come at the expense of scalability—the barcoding step is performed at the end, requiring one to work with multiple wells or plates, which can be more challenging for small-volume work. The BRB-Seq platform features an early barcoding step, during the reverse transcription priming with oligo-dT, allowing all samples to be multiplexed and enabling all work for hundreds of samples to occur in a single tube. This significantly simplifies the workflow; however, this is at the cost of decreased sensitivity for SARS-CoV-2 detection. It likely would benefit from a more robust reverse transcription priming step that isn’t reliant on the poly-A tail of mRNAs. As molecular technology matures, we believe that the existing trade-off between scalability and accuracy will decrease and allow for both scale and accurate detection across multiple pathogens.

### Limitations of our study

Our study, while the most comprehensive published study that directly compares multiple extraction kits and library preparation reagents, did not comprehensively survey all available kits and methods. We aimed to focus on full-service kits that were amenable to automation. This choice was due to the increasing use of automation in these labor-intensive assays across clinical laboratories.

We are only as good as the tools available for measuring the quantity of RNA. If the accuracy of our quantitation and RINe score is poorer or artificially low in these samples, it will lead to a higher starting RNA material. Second, given that the vast majority of the classified RNAs were non viral, it is most likely that the fragmented and degraded eukaryotic and bacterial RNA contributed to the low RINe scores. Similarly, the concentration of RNA is more heavily influenced by the amount of dominant human and bacterial RNA than by viral RNA. While we observed low overall SARS-CoV-2 genome coverage in our SARS-CoV-2 positive samples, we saw several non-overlapping regions of coverage in the Revelo and NEB libraries as well as reads tiling the expected region of coverage in the BRB-Seq libraries. It may be that additional optimizations must be conducted to capture an entirely intact viral genome from highly degraded sample types such as saliva, however the results are confounded by the source of the SARS-CoV-2, which was exogenous heat-inactivated SARS-CoV-2. It may be possible to have high-quality viral RNA masked by the much greater quantity of low-quality human RNA when looking at concentration and RINe score as quality metrics, however, further investigation with clinical samples should be conducted before making any such claims. There are no measures that can distinguish between human and non-human nucleic acids, but such a measure could be a valuable metric for agnostic infectious disease diagnostics.

Additionally, we tested the accuracy of SARS-CoV-2 detection across 24 platform combinations but did not conduct larger clinical validations due to cost and scale constraints. From this work, one would likely narrow down the combinations for further validation studies, which can accrue significant costs. While this would be a valuable study required for clinical implementation of any mNGS-diagnostic, we felt it was beyond the scope of this study and relevant for follow-up studies after determining the best-performing combination. We hope that our data can highlight the critical importance of identifying the best extraction and library preparation method for one’s intended purpose. While no single combination was superior across all metrics, we found that using a full-transcriptomic library preparation kit could compensate for poor sample quality.

### Optimizations to Improve Pathogen Detection

We considered several approaches that may improve pathogen detection. These methods, commonly used in human gene expression studies, focus on removing highly abundant reads, such as ribosomal RNA or hemoglobin from whole blood samples. The logic is that removing reads that represent an estimated 70-80% of all reads will allow more lowly expressed transcripts to be reverse-transcribed, amplified, sequenced, and detected. We observed improved detection of our spiked-in SARS-CoV-2 virus with the Revelo kit which utilizes SPIAboost to suppress human rRNA amplification. Alternative methods to deplete abundant reads include bead-based rRNA depletion^21^ and CRISPR-based approaches^22^. We did not include these additional depletion approaches in these studies as they increased the complexity of workflows, risked losing low-abundance reads through dropout^23^, and limited our ability for automation. Additionally, we chose to omit the optional addition of DNase to simplify our workflow and facilitate automation.

### Saliva specimens can provide high-quality data for clinical diagnostics

Although skepticism exists about using RNA extracted from saliva samples for clinical diagnostics^24^, our study demonstrates that we can extract nucleic acids of sufficient quantity and quality to generate diagnostic sequencing data compatible with clinical infectious disease testing workflows^9,25,26^. We found that samples extracted using a range of extraction kits yielded low but detectable RINe scores, and variable quantities of nucleic acid. However, the highest quality did not have a direct correlation with the highest detected spiked-in virus, MS2 and SARS-CoV-2. Samples that showed minimal RNA material on both TapeStation and Qubit assays could be used to produce high-complexity RNA-Seq libraries. Extracted nucleic acid with a low RINe score may produce an uninterpretable RNA-Seq library for differential mRNA expression analysis. However, based on our data, a low RINe score should not be used to disqualify a sample from being processed for mNGS viral pathogen detection.

As sequencing technologies advance, costs decrease, and high-throughput sequencing platforms become more widely available, it is crucial to identify which nucleic acid extraction methods enable the generation of high-quality sequencing libraries from imperfect clinical samples. We found that the Zymo Quick-RNA Magbead extraction method preserves the highest number of reads mapping to SARS-CoV-2 and MS2 phage RNAs across the full transcriptome library preparation methods. Among the RNA-Seq library preparation methods, we found that the Revelo RNA-Seq High Sensitivity library preparation kit captured the highest percentage of SARS-CoV-2 and MS2 phage reads across all extraction methods, albeit with high rates of duplicated reads. The choice of RNA-Seq library preparation kits can significantly alter the sensitivity of the assay. Added internal controls, such as MS2, could assist in determining RNA-Seq library quality post-library creation and sequencing, similar to MS2’s role in RT-PCR; however, a suitable quality metric has not yet been found that can be assessed before library creation. With a better understanding of the link between extraction quality and library preparation, our work highlights the need for quantitative measures of nucleic acid that better predict the success of RNA-Seq library creation in a mNGS context.

## DECLARATION OF INTERESTS

The authors declare no competing interests.

## ACKNOWLEDGEMENTS

This work was supported by R01AI177859 to VA and EE, NIGMS T32 GM008042 (2022-2024) and NIGMS T32 GM152342 (2024-2025) to KQ and CDC Grant 75D30123C18049. We thank the members of the Arboleda lab for their helpful comments and feedback on the manuscript. Icons in Figure 1 were partially provided by Servier https://smart.servier.com/ is licensed under CC-BY 3.0 Unported https://creativecommons.org/licenses/by/3.0/

## Author Contributions

E.E. and V.A.A. obtained funding for the project. V.A.A., E.E. and K.Q. designed the project. K.Q. performed the analyses. K.Q., V.A., D.K., and P.Z. performed the experiments to generate the data. T.T. assisted with obtaining the remnant samples.

## Figures Acknowledgements

Icons in Figure 1 were partially provided by Servier https://smart.servier.com/ is licensed under CC-BY 3.0 Unported https://creativecommons.org/licenses/by/3.0/

## Notes

### Competing Interest Statement

The authors have declared no competing interest.

